# Multi-omic profiling of Plasmodium knowlesi blood stages identifies host adaptation barriers

**DOI:** 10.64898/2026.10.02.756168

**Authors:** Samuel Pazicky, Nicolas Aranciaga, Xiahong Gao, Angjing Wen, Kotchanipa Rukruam, Seth Tjia, Huy Duong, Ka Diam Go, Pablo Bifani, Michal Kucharski, Peter Rainer Preiser, Zbynek Bozdech

## Abstract

The zoonotic parasite Plasmodium knowlesi has emerged as a widespread cause of human malaria in Southeast Asia. We assembled a high-resolution transcriptome of zoonotic *P. knowlesi* across its intraerythrocytic developmental cycle and combined transcriptomics, quantitative proteomics and thermal stability measurements to compare parasites cultured in macaque and human red blood cells. We found that *P. knowlesi* accelerates through the ring stage to settle at the same developmental pace as *P. falciparum* during the trophozoite and schizont stages. Subsequent multi-omic analysis revealed broad transcriptional and proteomic adaptations of *P. knowlesi* to the human host cell environment. Parasites cultured in human cells expressed glutathione synthetase earlier and displayed changes in upstream processes that include NADPH production by the pentose phosphate pathway and glutamine synthesis. These changes were accompanied by further differences in the expression of ribosomal subunits and mitochondrial respiratory complexes. Together, our findings offer a systems-level view of non-genetic adaptations of *P. knowlesi* to the human red blood cell environment.

## Introduction

Malaria is a major social and economic burden in equatorial countries, causing almost 300 million cases and 600 thousand deaths worldwide^1^. Although the majority of research focuses on the most lethal malaria parasite, *Plasmodium falciparum*, *P. vivax* makes up for two thirds of all malaria cases in South America and a zoonotic species, *P. knowlesi*, accounts for the majority of cases in some countries in Southeast Asia^1^. *P. knowlesi* is phylogenetically closely related to *P. vivax*: 80% of its genes have orthologs in *P. vivax*^2–4^, the invasion machinery of the two species is similar^5,6^ , and both of them prefer to invade reticulocytes^7^. Having been successfully adapted for continuous *in vitro* propagation^8^ and being readily amenable to genetic manipulation by Crispr/Cas9^9^, *P. knowlesi* became a model organism for *P. vivax* research. This allowed generating transgenic lines for studies of resistance-related genes^10^, production of antibodies^11^ , and their testing^12^. Recently, two large mutagenesis studies used the piggyBac transposon system to establish the essentialome of *P. knowlesi*^13,14^.

However, *P. knowlesi* itself has recently emerged as the dominant human-infecting malaria species in Southeast Asia. Previously mistaken for microscopically similar *P. malariae,* human *P. knowlesi* infections were first recognized in Malaysian Borneo in 2004^15^ and later found to be prevalent across entire Malaysia and potentially lethal^16^. Although the fatal *P. knowlesi* cases display similar pathology to *P. falciparum*^17^, a higher proportion of *P. knowlesi* malaria patients develop severe malaria^18–22^. Although early medical intervention can reduce the case fatality rates^23^, *P. knowlesi* has become a dominant human malaria species in some parts of Southeast Asia where *P. falciparum* is approaching elimination^24^. Indeed, 5.1-7.1% of the population surveyed in distinct sites in Malaysia and the Philippines displayed *P. knowlesi* seropositivity^25,26^.

Despite these numbers, it remains elusive whether all *P. knowlesi* infections have zoonotic origin. The natural hosts of *P. knowlesi* are Old World macaques, with the long-tailed macaque *Macaca fascicularis* serving as the main parasite reservoir in Southeast Asia^27^. The macaque-infecting parasite population maintains higher genetic diversity than in human infections, which has been suggested to sustain a baseline level of human cases via spillover from the reservoir^28^. Indeed, modelling individual case reproduction numbers from a decade of Malaysian surveillance data found *P. knowlesi* transmission to be consistently subcritical for sustained human-to-human transmission^29^. This is supported by the fact that engagement in agriculture and forest-near activities are the highest risk factors for *P. knowlesi* infection and the strongest predictors of seroconversion^25,30^. Nevertheless, direct human-to-human transmission via mosquitoes has been demonstrated on human volunteers in laboratory settings^31^. *P. knowlesi* gametocytes have been detected in 47% and 67% of patients surveyed in Vietnam and Malaysia, respectively^32,33^ , and mosquitoes fed on cultured *knowlesi* gametocytes grown in human red blood cells (RBCs) as well as blood from *P. knowlesi*-infected patients developed oocysts^34,35^.

While the adaptation mechanism for invasion of human RBCs via normocyte-binding protein Xa has been established^36^, the subsequent zoonotic adaptation, prerequisite to the production of viable gametocytes, has not been investigated so far. Here, we established a high-resolution multi-omics dataset capturing the transcriptional, proteomic, and metabolomic profiles of the zoonotic *P. knowlesi* intracellular developmental cycle (IDC) in human RBCs. Comparing these to *P. knowlesi* cultured in macaque RBCs, we revealed specific signatures of transcriptional, proteomic, and metabolic reprogramming that are specific to the zoonotic adaptation processes. Specifically, *P. knowlesi* rewires glutamine biosynthesis and pentose phosphate pathways to presumably regulate the production of glutathione that compensates for the higher oxidative environment in human RBCs. This is accompanied by dysregulation of mitochondrial respiratory complexes and ribosomal integrity in human RBCs, where the parasites increase production of ribosomal mRNAs but degrade the ribosomal proteins and the associated 18S rRNA.

## Results

### Transcriptional time course of *P. knowlesi* IDC

*Plasmodium* species are characterized by just-in-time gene expression that results in regular oscillations of mRNA levels along the IDC^37^. In this transcriptional cascade, functionally related and physically interacting proteins are transcriptionally co-regulated, which constitutes one of the key foundational principles of the malaria parasite asexual blood stage development. To investigate this in *P. knowlesi,* first we wished to reconstruct the high-resolution genome-wide IDC transcriptome map. For this, we cultured a highly synchronized *P. knowlesi* A1H1 parasite strain in human red blood cells (hRBCs) and collected samples across its IDC: 0–34 hours post-invasion (hpi) at 2-hour (h) intervals (Fig. 1a). To generate the IDC transcriptome map, we subsequently isolated total RNA fractions from each of the samples and subjected them to RNA-Seq analysis as previously described^38^ (see Methods). Overall, we detected 5,465 genes (5,183 without rRNA and var genes) in two highly correlated biological replicates (Pearson’s r=0.89, Fig. 1b, Extended Data Fig. 1a, Supplementary Data 1 & 2). The principal component analysis (PCA) analysis confirmed a high inter-replicate consistency as well as the expected cyclical IDC trajectory in which we estimated completion of the cycle from the initial state (0h) in 31 hours, reflecting the duration of the IDC (Fig. 1c, Extended Data Fig. 1b). To quantify the periodic oscillations, the expression values were fitted with periodic function that in turn allowed estimations of the peak expression time for each gene. These were consistent (median phase difference Δφ=0h) and exhibited tight synchronicity (median |Δφ|=1.6h, Fig. 1d). Replicates displayed a good peak correlation across all genes (intraclass correlation ICC=0.73, Fig. 1e), which was even stronger for genes with a clear periodic transcriptomic profile (R^2^>0.5 ICC=0.93). Given this consistency of the transcriptomics results, we merged the two biological replicates to build the consensus IDC transcriptome of the *P. knowlesi* parasites (Fig. 1f). Gene set enrichment analysis (GSEA) at peak expression times (FDR<0.05) revealed a functional temporal pattern, with expression of genes encoding Kelch13-interacting proteins and factors of RNA processing and translation peaking in rings; expression of genes related to metabolic processes and mitochondria peaking in trophozoite; genes involved in the processes and complexes in DNA replication and cell division exhibiting expression peaks in late trophozoites or early schizonts; and genes in organelles and complexes with functions in parasite egress and invasion finally reaching the expression peaks in mature schizonts (Fig. 1g-h, Supplementary Data 3). Overall, this temporal pattern of metabolic and cellular functions in *P. knowlesi* resembles that of *P. falciparum*, suggesting that the basic developmental program is conserved between the *Plasmodium* species. This basic developmental conservation was demonstrated in our earlier study across six human and rodent *Plasmodium* species^39^. Here we, however, also demonstrated divergences in some elements of the IDC transcriptional cascades that presumably reflect specificities of each *Plasmodium* species, determining the adaptations to their host environment and infection niches.

**Figure 1.**
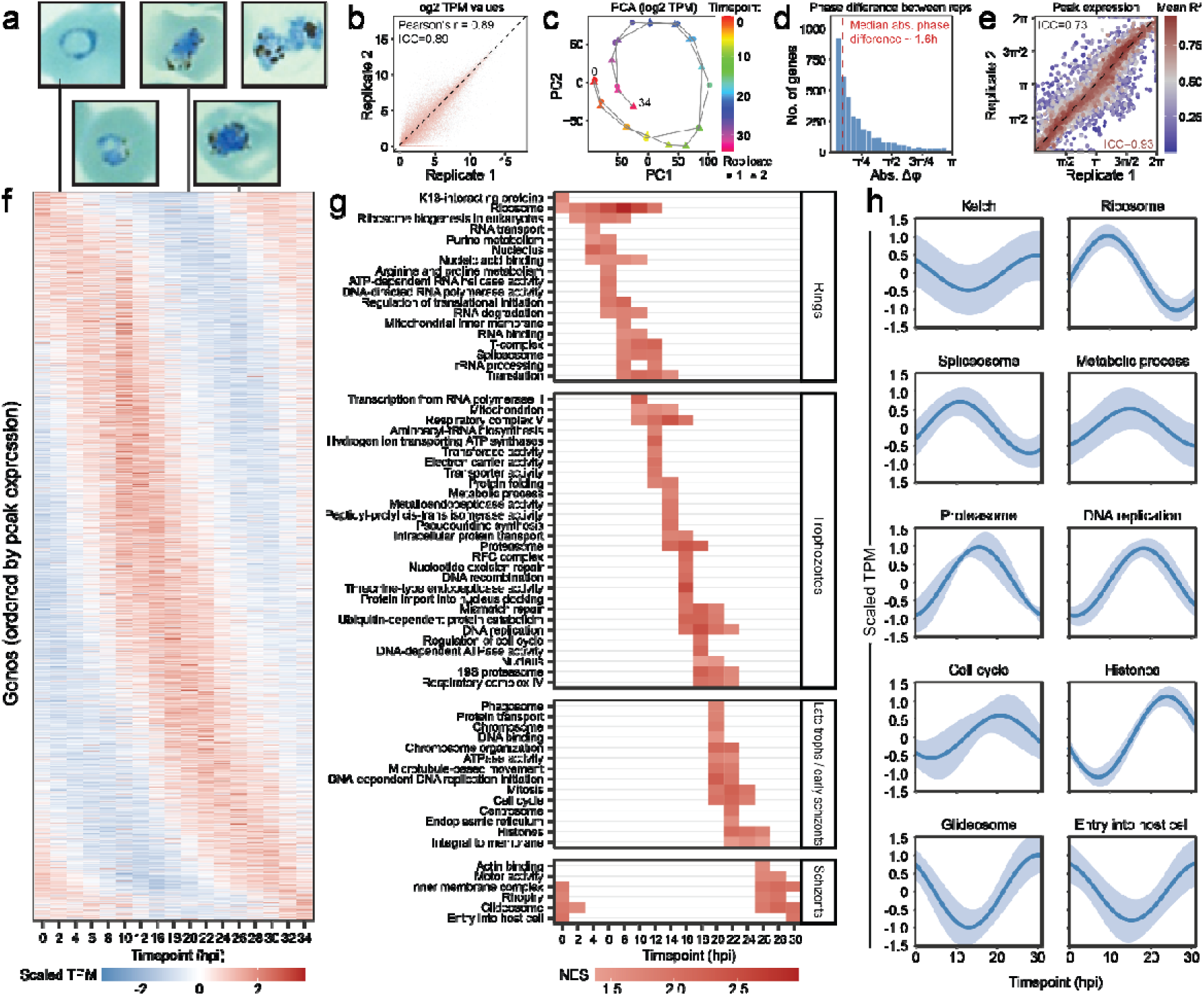
2h transcriptional time course of P. knowlesi IDC in human RBCs. **a** Representative Giemsa-stained parasites at 2, 8, 20, 26 and 32 hpi. **b** Scatter plot of log2 TPM values from all timepoints between two replicates shows excellent agreement (Pearson’s r=0.89, ICC=0.89). **c** PCA based on log2 TPM values matched shows a good agreement between the replicates. **d** Histogram of absolute phase difference values between two replicates shows that majority of genes showed synchronized expression profiles within ±1.6h. **e** Scatter plot of peak expression times of the two replicates shows good agreement (ICC=0.73 but for profiles fitted with R^2^>0.5 ICC=0.93). The less correlated genes (not on diagonal) often display low R^2^ values (blue points), suggesting that the expression of these genes could not be described by cosine fit. **f** Heatmap of z-scaled consensus TPM values of *P. knowlesi* genes. **g** Gene set enrichment analysis at every timepoint shows timed expression of functionally related genes and complex subunits. **h** Fitted z-scaled consensus TPM values of chosen enriched terms with confidence intervals.

### P. knowlesi compresses ring-stage development to align with P. falciparum

To explore this for *P. knowlesi* grown in the zoonotic, heterologous host of human erythrocytes, we compared the expression patterns of orthologous genes between *P. knowlesi* consensus IDC and previously published data from *P. falciparum in vitro* and *P. vivax ex vivo* cultures^40,41^. Specifically, we generated scatterplots of IDC peak expression timing of the three human-infecting *Plasmodium* species and fitted both linear regression trendlines (Fig. 2a & b). Interestingly, the peak expression trend between *P. knowlesi* and *P. falciparum* was characterized by two segments: first (0-10.9 hpi), in which *P. knowlesi* appeared to progress ∼2.5x faster than *P. falciparum*, and second (10.9-31 hpi), in which both species developed approximately at the same rate (Fig. 2a & b). In contrast, the trend between *P. knowlesi* and *P. vivax* gene expression patterns appeared to be linear throughout the entire IDC with a constant relative developmental rate (1.5x faster, Fig. 2b). These marked differences in the two comparative patterns of gene expression were confirmed by the analysis of interspecies phase differences fitted with a saturating exponential model (Fig. 2c & d). The results show that *P. knowlesi* reached the expression difference of 17h (48h *P. falciparum* – 31h *P. knowlesi* IDC) after 11 hpi and finished the last 20 hours of the IDC at the same speed as *P. falciparum* (Fig. 2c). In contrast, the phase difference linearly increased to 17h when compared to *P. vivax,* showing that the pacing of *P. knowlesi* and *P. vivax* development is comparable across the entire IDC (Fig. 2c). Next, to evaluate the functional differences between *P. falciparum* and *P. knowlesi* IDC programmes, we first used the two-segment fit to adjust *P. falciparum* expression times in order to remove the global differences caused by different IDC pacing (Extended Data Fig. 2a). After the adjustment, both *P. falciparum* and *P. vivax* show good temporal consistency with *P. knowlesi* (Pearson’s r = 0.74 and 0.77, Extended Data Fig. 2a-c). Compared to *P. falciparum*, the shifts were larger, reaching half of the cycle for cell surface proteins (Fig. 2e, Extended Data Fig. 2d). Gene sets associated with mitochondrion, respiration, TCA cycle, and rRNA processing were expressed earlier in *P. knowlesi*, while the glideosome, 19S proteasome, spliceosome, and genes associated with mitosis displayed delayed expression (Supplementary Data 4). Comparison to *P. vivax* showed only small shifts (±4h), some of which were identical to those from *P. falciparum* comparisons. These included genes related to respiration, ribosomes, and the 19S proteasome, which are likely related to the specifics of *P. knowlesi* biology. Uniquely to the *P. vivax* comparison, certain metabolic pathways and ribosomal proteins were expressed earlier in *P. knowlesi* (Fig. 2f, Extended Data Fig. 2e, Supplementary Data 4). This suggests that the observed temporal divergence in *P. knowlesi* gene expression has a functional relevance, presumably shifting key metabolic and cellular functions according to its needs when developing in human erythrocytes. Overall, zoonotic *P. knowlesi* displays larger IDC progression differences compared to *P. falciparum* and a “time compression” of the ring stage such that the transcriptional profile of the later stages (trophozoites and schizonts) aligns with the last 20 hours of *P. falciparum* development. On the other hand, *P. vivax* and *P. knowlesi* display similar progression in the IDC transcriptional cascade, indicating closer resemblance in their asexual blood stage development.

**Figure 2.**
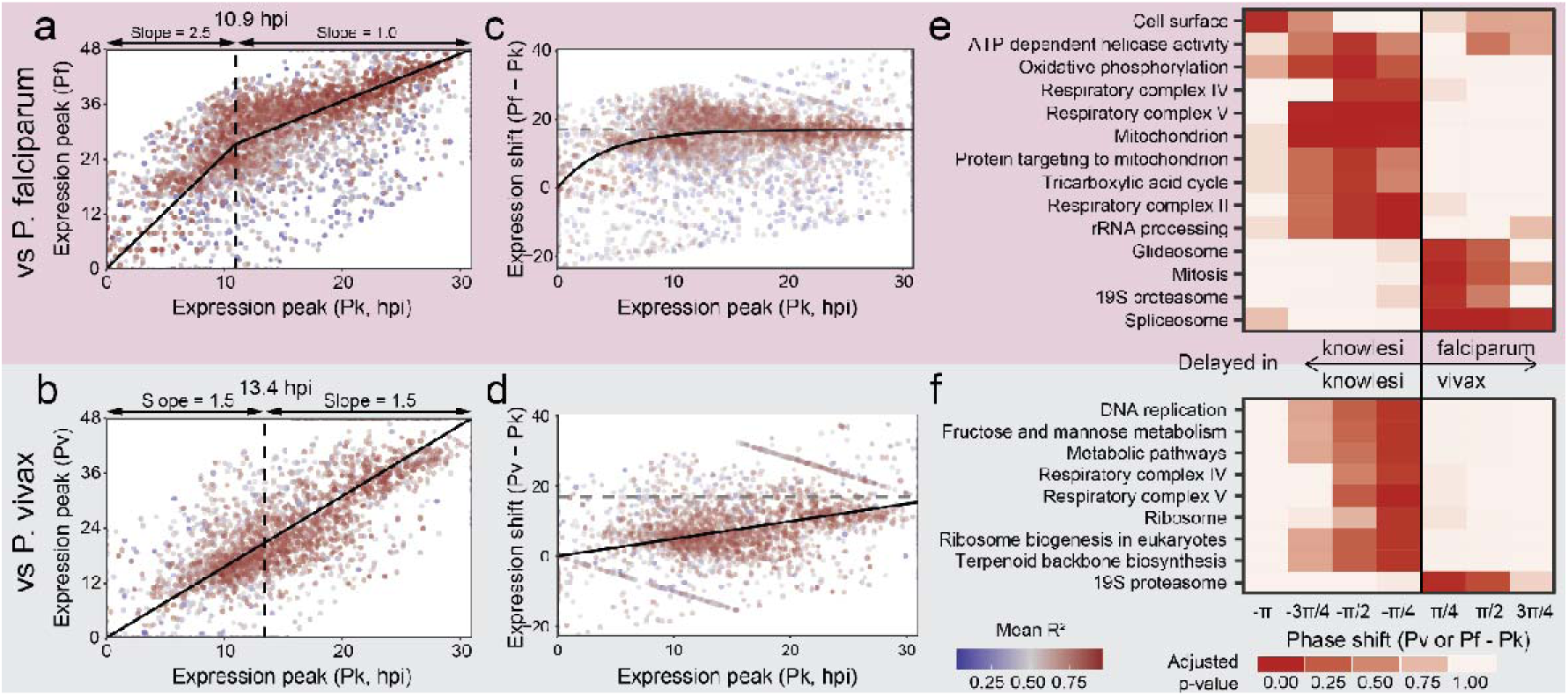
*P. knowlesi* rushes through the ring stage compared to *P. falciparum*. **a** Scatter plot of *P. knowlesi* and *P. falciparum* peak expression times and two-segment linear fit splits the *P. knowlesi* IDC into fast-paced ring stage (2.5x the speed of *P. falciparum*) and trophozoite/schizont stage that progresses at the same speed as *P. falciparum*. **b** Correlation between gene expression peak times of *P. knowlesi* and circular phase differences between *P. falciparum* and *knowlesi* shows identical split between ring and trophozoite & schizont stages. **c,d** The same mathematical analyses for comparison of *P. knowlesi* with *vivax* show that the pacing of their IDCs is identical across all IDC stages with constantly 1.5x faster *P. knowlesi* parasites. **e,f** GSEA across a series of phase shift between *P. knowlesi* and adjusted *P. falciparum* or *P. vivax* peak expression times shows shared and species-specific differences.

### *P. knowlesi* shows host-dependent differences in gene expression

In the next step, we wished to compare the IDC transcriptional cascade of *P. knowlesi* developing in its zoonotic (human) host to that in the natural (macaque) host. For that, we cultivated highly synchronized *P. knowlesi* parasites in hRBCs and RBCs purified from blood of long-tail *Macaca fascicularis* macaque (mRBCs). Using the identical RNA-Seq approach^38^ (above), we reconstructed the IDC transcriptional cascades in both hosts in two replicates (Fig. 3, Extended Data Fig. 3a). To compensate for small differences in the parasite growth rates, we first aligned the gene expression data to the consensus hRBC IDC (Fig. 1f) and re-calculated the timepoints correspondingly (Extended Data Fig. 3b). Both replicates displayed good correlation between human and macaque gene expression peak times (Pearson’s r=0.87 and 0.74, Fig. 3a and Extended Data Fig. 3c) and displayed tight inter-species synchronicity (median |Δφ|=2.5h for both replicates, Fig. 3b). Interestingly, both mRBC- and hRBC-cultured parasites displayed the compressed ring stage compared to *P. falciparum,* which suggests that this feature of the IDC is not impacted by zoonotic growth (Extended Data Fig. 2f-g). Nevertheless, we observed considerable fluctuations in the temporal gene expression pattern between the *P. knowlesi* IDC within the zoonotic (hRBCs) compared to the natural (mRBC) host, with temporal shifts within the Δφ< /4 interval, including a set of 18 genes that exhibited dramatic shifts, with their peak expression offset by Δφ> /4 (Fig. 3c & d, Supplementary Data 5).

**Figure 3.**
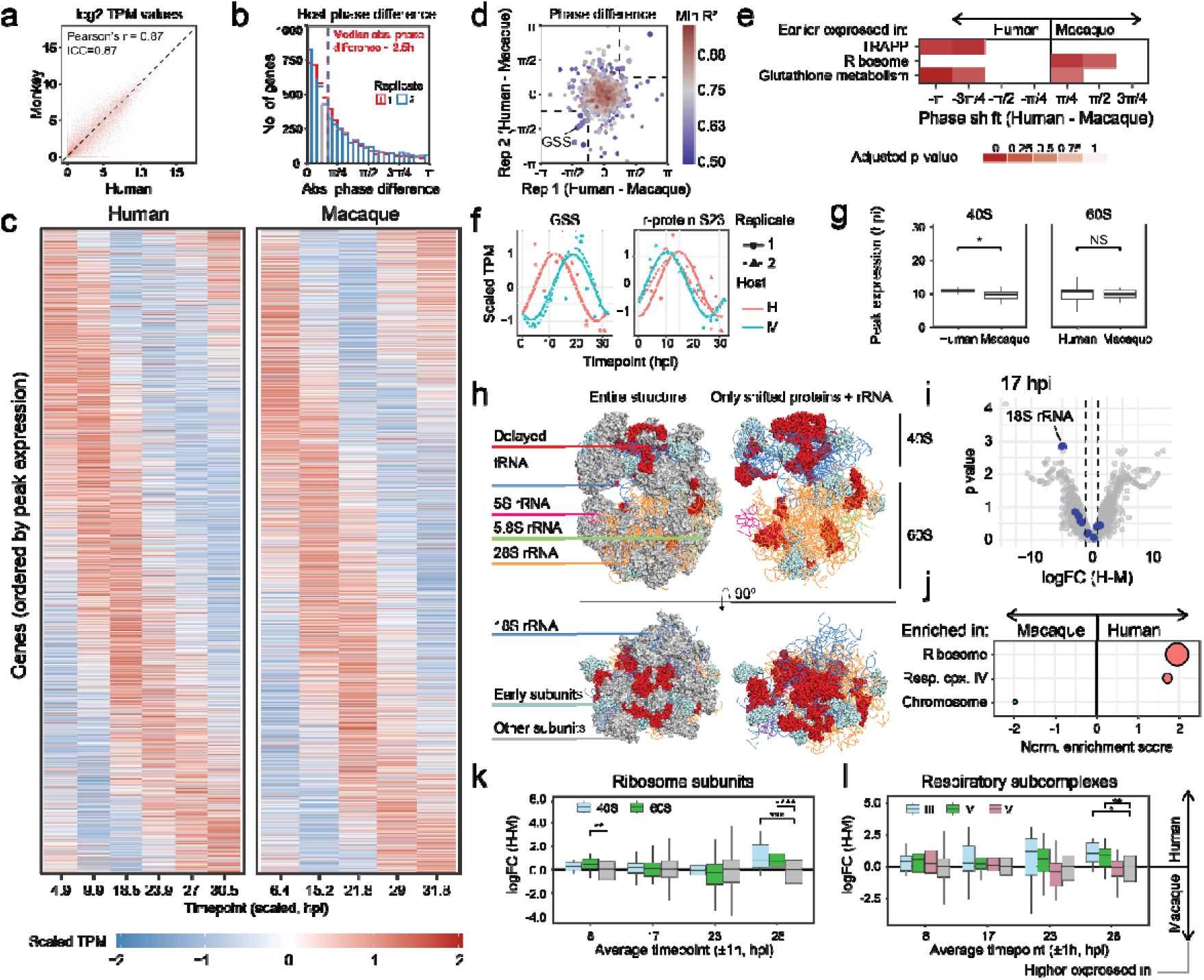
Differences in transcription time course between human and macaque host. **a** Scaled TPM values in human and macaque host with genes ordered by peak expression (replicate 2). **b** Histogram of absolute phase difference values between two host RBCs shows that majority of genes showed synchronized expression profiles within ±2.5h. **c** Phase difference between peak expression of genes in human and macaque RBCs in replicate 1 and 2. GSS=glutathione synthetase. **d** Heatmap of gene expression profiles in human and macaque RBCs, ordered by average peak expression time. **e** Heatmap colored by p-values from one-sided GSEA analysis shows differences in expression timing of groups of genes. **f** Fitted gene expression profiles of GSS and ribosomal subunit S23. **g** Difference in expression peak of 40S and 60S ribosomal subunits in human and macaque RBCs (Wilcoxon test). **h** Structure of the ribosome (PDB 9j9h) with (left) or without (right) proteins that did not have shifted gene expression between human and macaque RBCs. Proteins colored in red were delayed in hRBCs. **i** Differential expression analysis at 17 hpi show 18S rRNA as one of the most downregulated genes. **j** GSEA based on logFC values from comparing gene expression values between human and macaque RBCs at 28 hpi. **k-l** Comparison of distribution of host-differences of all genes (gray) and either 40S and 60S ribosome subunits (**k**) or respiratory complexes III, IV and V (**l**). Wilcoxon test.

GSEA across a series of phase-differences revealed three enriched terms (FDR < 0.1, Supplementary Data 6), including Transport protein particle (TRAPP) complex and glutathione metabolism exhibiting earlier and ribosome structure with later onset of their expression in the zoonotic versus the natural host (Fig. 3e). Specifically, genes involved in glutathione metabolism peaked almost half of IDC earlier in hRBCs, which was predominantly driven by glutathione synthetase (GSS, PKA1H_010015900), with peak expression at 12.2 hpi in hRBCs compared to 19.2 hpi in mRBCs (Fig. 3f). As the terminal enzyme in the glutathione synthesis that is essential in protecting the parasite from a highly oxidative environment (reviewed by Müller et al^42^), this temporal shift in expression may indicate its particular role in the *P. knowlesi* zoonosis (see below). TRAPP is a membrane-tethering complex that mediates vesicular transport between organelles^43^ that has not been investigated in *Plasmodium* species so far. Interestingly, the enrichment was driven by the TRAPPC4 subunit (Extended Data Fig. 3d) that is additionally implicated in cell signalling and gene expression regulation in cancer cells^44,45^. The gene expression delays of the ribosomal structure appeared to be driven entirely by the 40S subunit genes (p<0.05), while no significant temporal shift in the expression could be detected for several 60S subunits (Fig. 3f&g). To gain mechanistic insights, we mapped all ribosomal genes with shifted (±1.5 h) expression profiles onto the structure of the *P. falciparum* ribosome^46^ and found that 9 out of the 13 proteins encoded by hRBC-delayed genes directly interact with 18S rRNA (Fig. 3h). We hypothesized that the shift in ribosomal gene expression could be associated with changes in 18S rRNA levels. To investigate overall differences in RNA levels between hRBCs and mRBCs, we performed differential gene expression analysis at four paired time points (8, 17, 23, and 28 hpi, ±2 h, Supplementary Data 7). Indeed, 18S rRNA was among the most significantly downregulated transcripts in hRBCs at 8 and 17 hpi (Fig. 3i, Extended Data Fig. 3e, Supplementary Data 8). Inspecting the datasets from the individual replicates, 18S rRNA was downregulated in 7/8 total time points compared to other rRNAs (Extended Data Fig. 3f).

To complement the functional analysis based on temporal shifts (Fig. 3e), we next performed GSEA based on overall differential expression and found that only the schizont timepoint contained significantly enriched gene sets, which, accordingly, include the ribosome being overexpressed in hRBCs (Fig. 3j, Supplementary Data 9). Inspecting the change distribution of 40S and 60S subunit genes across the four timepoints, there is ∼2-fold overexpression of both subunits at 28 hpi (schizonts) in hRBCs compared to mRBC (Fig. 3k). In addition to the ribosome, early expression of mitochondrial respiratory complexes in hRBC-resident parasites was just under the significance threshold in the expression shift GSEA and plotting their fold change distributions, we found complexes III and IV to be enriched in hRBC-cultured parasites starting at 23 hpi and significantly enriched at 28 hpi (Fig. 3l). Interestingly, respiratory complexes IV, and V were also expressed earlier in hRBC-resident *P. knowlesi* compared to both *P. falciparum* and *P. knowlesi* (Fig. 2e-f). This transcriptional analysis revealed significant shifts in transcription of few but potentially essential stress-as well as growth-related pathways, including glutathione synthesis, ribosomal structure, and mitochondrial complexes, that could reflect the specific needs of *P. knowlesi* to survive in its zoonotic host during the IDC.

### Host-specific parasite proteomes revealed shifts in the pentose phosphate pathway for NADPH for glutathione regeneration in the zoonotic hRBCs host

Gene expression changes can alter protein abundances, but their effects are often attenuated or even reversed by additional layers of regulation during protein translation and degradation^47^. Hence, we carried out quantitative proteomics analysis to complement the observed transcriptional differences between *P. knowlesi* in human and macaque RBCs. For that, we sampled four time points of the parasite’s IDC in the two hosts and measured their protein abundances by mass spectrometry. Reflecting the cyclical regulation of the gene expression, the abundances of 1,570 detected parasite proteins also followed periodic abundance profile (Fig. 4a, Supplementary Data 10) that displayed somewhat more variation compared to the gene expression profiles (median |Δφ|=3.9h vs 2.5h), possibly reflecting additional layers of posttranscriptional regulation (Fig. 4b). The delay of protein abundance peaks after the gene expression peaks corresponded to 11-16% of the IDC (median 5.0h and 3.5h in mRBCs and hRBCs, respectively, Fig. 4c), pointing to a faster pace of translation in *P. knowlesi* compared to *P. falciparum* (33% of *P. falciparum* IDC^48^. Finally, we found that the IDC driven oscillations in gene expressions were attenuated 4-5x at the protein levels compared to RNA level (median amplitudes 1.4 vs 5.6 or 6.8, Fig. 4d). The key example is GSS that we found to have a temporal shift at the RNA level (4-7-fold across the IDC) that was also reflected by the protein abundance shifts (1.1-1.2-fold) (Extended Data Fig. 4a). Importantly, the premature gene expression of GSS in hRBCs (Fig. 3f) was mirrored at the protein level (peak at 18.1 vs 26.6 hpi, Fig. 4e) with 5.9 and 6.9h gap between RNA and protein expression peaks in hRBCs and mRBCs, respectively.

**Figure 4.**
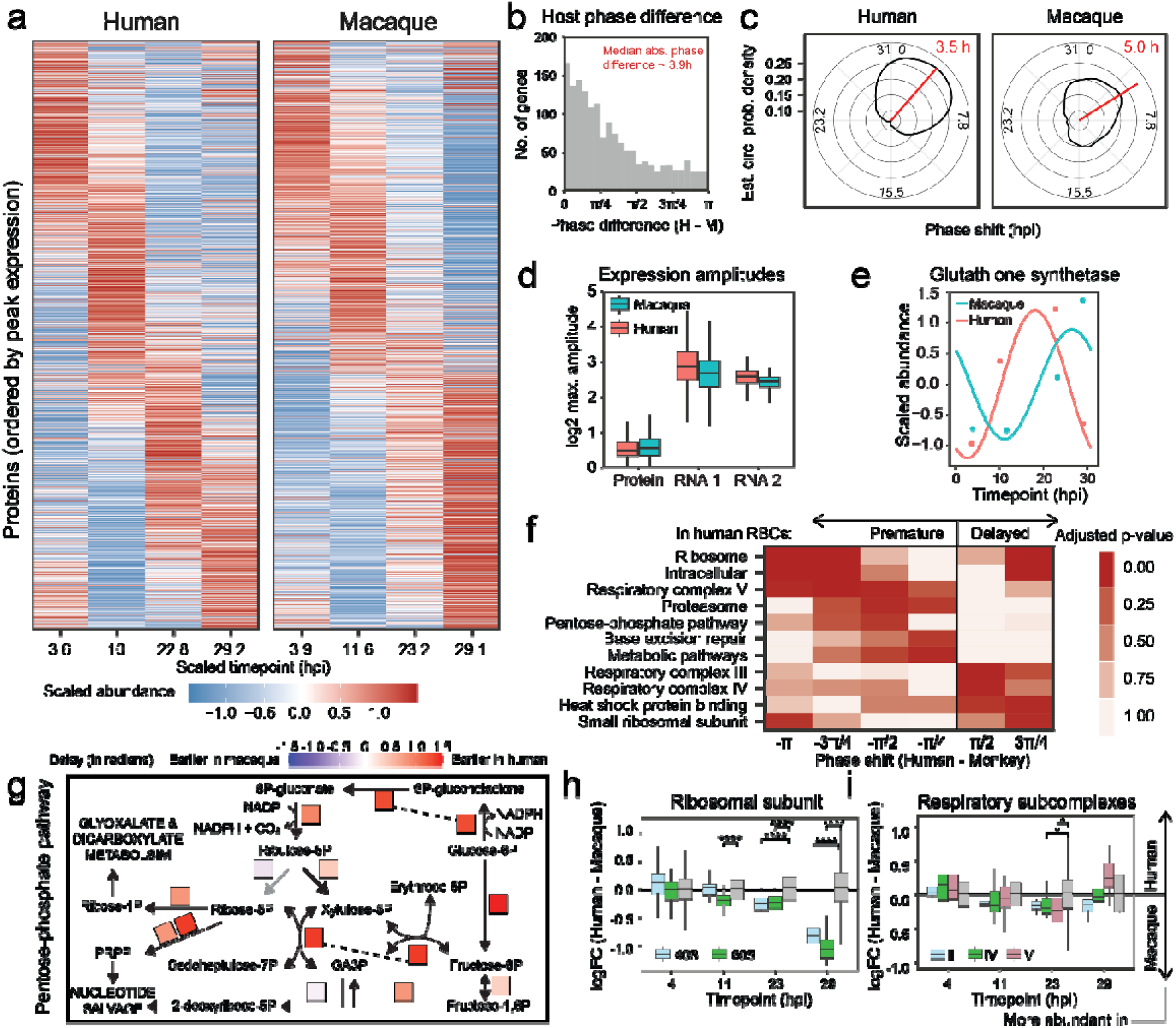
Differential time course proteomic analysis of *P. knowlesi* in human and macaque host cells. **a** Heatmap shows relative synchronicity of protein abundance oscillations between human and macaque RBCs. **b** Histogram of absolute phase differences shows larger differences compared to transcriptomic data (3.9 vs 2.5h). **c** Circular probability densities show that the delay of protein expression after RNA expression is shorter in human cells than macaque cells. **d** Amplitudes of gene expression oscillation are much larger compared to the oscillation of protein abundances. **e** Fitted protein abundance oscillation of glutathione synthetase in human and macaque RBCs show differential temporal regulation. **f** GSEA analysis across phase shifts identifies groups of proteins with different protein abundance peaks between human and macaque RBCs. **g** Pentose-phosphate pathway with squares representing the enzymes colored by delay of hRBC form mRBC expression. **h** Boxplots show that proteins of both ribosomal subunits decrease their abundance in later IDC stages in human RBCs compared to macaque RBCs. **i** Boxplots show that the protein abundances of subunits of three mitochondrial respiratory complexes (III, IV and V) decrease in the trophozoite stage in human compared to macaque RBCs. The p-value were calculated by Wilcoxon rank-sum test and adjusted by Benjamini-Hochberg.

To systematically assess the phase shifts at the proteome level, analogously to our observations at the RNA level, we performed GSEA and observed a broad spectrum of cellular and metabolic pathways expressed at earlier time points of the IDC in the zoonotically developing *P. knowlesi* compared to its natural host (Fig. 4f, Supplementary Data 11). These include the ribosome, proteasome, respiratory complex, base excision repair, and pentose phosphate pathway. In contrast, factors of the respiratory complexes III and IV, heat shock protein binding, and the small ribosomal subunit were significantly enriched amongst the genes whose protein abundance peaks were delayed in hRBCs. Notwithstanding that these shifts in protein levels were not completely reflected in the transcriptional shifts (Fig. 3), there are several biologically relevant links between the two levels of gene expression. First, the pentose phosphate pathway (PPP, Fig. 4g) is well established as the main source of NADPH that is essential for recycling of glutathione^49^. NADPH is produced by two PPP enzymes: glucose-6-phosphate dehydrogenase-6-phosphogluconolactonase (G6PD-PGLS, PKA1H_120033100) shifted the expression already in ring stages from 11.6h in hRBCs to 5.2h in mRBCs; and 6-phosphogluconate dehydrogenase (PKA1H_120032200) shifted the expression from 24.1h to 19.5h in hRBCs, in sync with GSS (Extended Data Fig. 4b-c). This early shift of G6PD-PGLS may indicate the need for a higher level of glutathione for the hRBC-resident *P. knowlesi* parasites, presumably facing higher oxidative stress early in the IDC. In future experiments, such an intriguing possibility could be further explored, possibly as a crucial aspect of *P. knowlesi* zoonosis.

Next, we focused on the protein-level temporal shift in the expression of the ribosomal subunits detected by the GSEA (Fig. 4f) as they also displayed temporal shift in mRNA abundance (Fig. 3). Examining the ribosome levels timepoint-by-timepoint, we found that the abundance difference is increasing along the IDC, with the biggest downregulation in hRBCs compared to mRBCs at the last timepoint (Fig. 4h). Broader GSEA analysis based on differential protein expression across all timepoints revealed additional differences in levels of proteins involved in RNA transport (driven by initiation factor 3 complex subunits) and T-complex subunits (Extended Data Fig. 4d & e). Initiation factor 3 (IF3) and T-complex act upstream and downstream of ribosome translation, respectively. Curiously, the protein abundance of the respiratory subcomplexes II, IV, and V was decreased in hRBCs compared to mRBCs from ring to trophozoite stages but recovered to the normal level (III and IV) or increased (V) in the schizonts (Fig. 4i). Taken together, the IDC development of *P. knowlesi* in its zoonotic host is marked by significant differences in levels and expression timing of proteins, involving key factors of the parasite growth and development ranging from protein metabolism through energy and redox metabolism to DNA repair. Although largely correlated, there were some marked discrepancies between the transcriptome- and proteome-based datasets. This likely reflects a complex relationship between transcription and posttranscriptional regulation during the *Plasmodium* IDC, which warrants further studies (see below).

### Parasite proteome exhibits conserved stability oscillation across the IDC

Thermal proteome profiling (TPP) has recently emerged as a new technique that, in addition to expression levels reflected by standard quantitative transcriptomics and proteomics, can closely reflect the functional state of the proteome^50–52^. In our recent work, we demonstrated the utility of TPP to study assemblies of protein complexes but also functions of entire pathways in the *Plasmodium* parasites^46,48^. Therefore, we performed thermal proteome profiling (TPP) in live parasites grown either in human or macaque RBCs (Fig. 5a), detecting between 803 and 2,644 proteins across eight samples (Extended Data Fig. 5a, Supplementary Data 12). The thermal stability of individual parasite proteins, as measured by melting temperature (Tm), was comparable between the parasites grown in hRBCs and mRBCs across the *P. knowlesi* IDC, (Pearson’s r=0.7-0.8, Extended Data Fig. 5b). Similar to abundance profiles, there were periodic oscillations of the protein stabilities across the IDC (median amplitude 2.5 and 3.8°C in hRBCs and mRBCs, Extended Data Fig. 5c) that generally correlated within the two hosts (median Pearson r=0.63, Extended Data Fig. 5d). We observed no correlation between the protein abundance and protein stability profiles, indicating that TPP provides an additional layer of information related to the protein functions (Extended Data Fig.5e). As such, it is feasible to expect that the stability differences could also reflect biological processes relevant to the *P. knowlesi* zoonotic adaptation.

**Figure 5.**
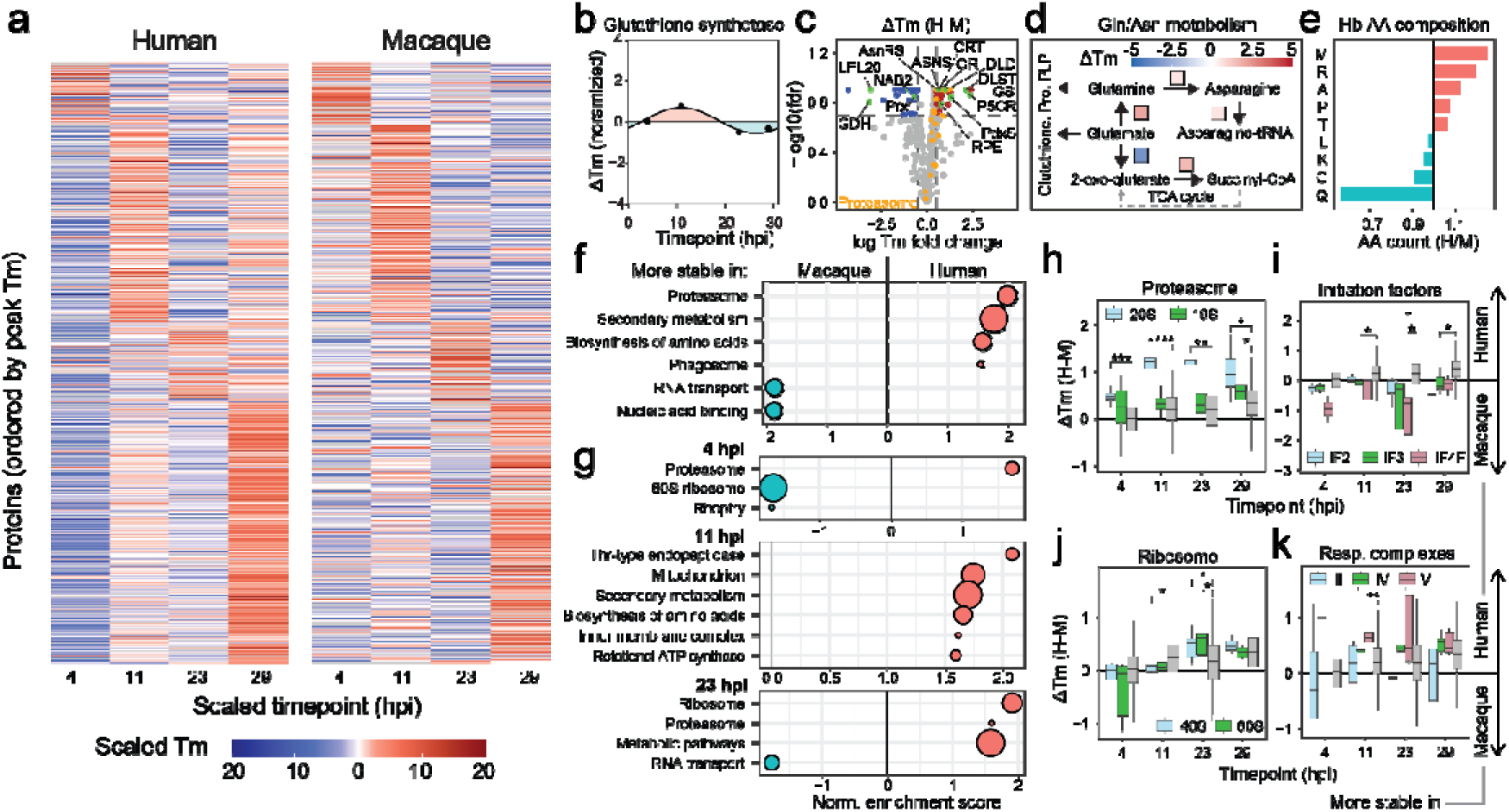
Oscillating stability changes provide functional insights into *P. knowlesi* proteome. **a** Heatmap showing conserved melting temperature (Tm) oscillation in human and macaque RBCs. **b** Glutathione synthetase exhibits period changes in stability with higher Tm in hRBCs at earlier stages and higher Tm in mRBCs at later stages. **c** Differential stability analysis shows significantly stabilized proteins in hRBCs or mRBCs. GDH – glutamate dehydrogenase; LPL20 – lysophospholipase 20; Prx – Peroxiredoxin; NAB2 – nuclear poly(A) binding protein 2; AsnRS – Asparagine-tRNA synthetase; ASNS – Asparagine synthetase; CRT – chloroquine resistance transporter; GR – glutathione reductase; DLD – Dihydrolipoyl dehydrogenase; DLST – dihydrolipoyl succinyltransferase; GS – glutamine synthetase; P5CR – Pyrroline-5-carboxylate reductase; PdxS – Pyridoxal phosphate synthase; RPE - Ribulose-phosphate 3-epimerase. **e** Ratio of amino acid residue composition between human and macaque hemoglobin shows that glutamine is the most underrepresented residues in human hemoglobin. **f** GSEA based on the differential stability. Positive values mean relative higher stability in hRBCs, negative in mRBCs. **g** GSEA of ΔTm values calculated separately for each timepoint. **h-k** Differences in normalized melting temperatures (ΔTm, hRBCs-mRBCs) of proteasome subunits (**h**) and initiation factors (**i**), ribosome subunits (**j**) and respiratory complexes (**k**) across four timepoints.

Indeed, GSS, which displayed consistently altered expression in hRBCs at mRNA (12.2 vs 19.2 hpi, Fig. 3b) and protein (18.1 vs 26.6 hpi, Fig. 4d), was also stabilized in hRBCs at an even earlier timepoint of the IDC compared to the mRBC (peak difference at 10.8 hpi, Fig. 5b, Supplementary Data 13). To expand on this observation, we calculated overall differences in Tm values (ΔTm) between the parasite proteins grown in the two host RBCs identifying additional proteins with differential thermal stability profiles including, glutathione reductase (GR) and ribulose-phosphate-3-epimerase (RPE) were stabilized in hRBCs across the entire IDC (Fig. 5c, Supplementary Data 14). Canonically, GR uses NADPH produced by PPP to reduce the oxidized glutathione and RPE acts in PPP just downstream of NADPH production and thus their thermal stability shifts likely related to that of GSS. Related to this, we also found large stability changes in enzymes involved in glutamate and glutamine metabolism. Glutamine synthetase (GS) and glutamate dehydrogenase (GDH), enzymes that synthesize either glutamine or 2-oxoglutarate from glutamate, were significantly stabilized and destabilized, respectively (Fig.5c & d). Enzymes that use glutamine as a substrate also had increased stability in hRBCs, including pyridoxal-5’-phosphate (PLP) synthase and asparagine synthetase (along with asparagine tRNA synthetase, Fig. 5c & d). Strikingly, glutamine is the most underrepresented amino acid in human α- and β-hemoglobin (4 residues in human vs 7 in macaque, Fig. 5e, Extended Data Fig. 5f).

For deeper insights into the host-differential proteome stability, we explored the stability differences GSEA (Fig. 5f, Extended Data Fig. 5g, Supplementary Data 15), which revealed generally increased protein catabolism in hRBCs, driven mainly by the 20S proteasome subunits and ubiquitin-like modifier HUB1 (<u>H</u>omologous to <u>Ub</u>iquitin). Additionally, the term phagosome is related to the parasite’s food vacuole, where the stabilized proteins are dominated by hemoglobin-digesting proteases (M17 leucyl aminopeptidase, falcilysin and knowpains 1 and 4) and include chloroquine resistance transporter (Fig. 5c). The only terms that showed higher stability in mRBCs than in hRBCs were the Nucleic acid binding and RNA transport that encompass mainly translation elongation and initiation factors. We found that this was driven mainly by the initiation factor 3 (IF3) and 4F (IF4F), and the largest effect was observed for subunit IF3α. IF3α is a classic example of a moonlighting protein^53^. We used MAP-X to profile the moonlighting in *P. knowlesi* proteome-wide and found 28 and 19 moonlighters in hRBCs and mRBCs of which 12, including IF3α, were shared, confirming our previous findings from *P. falciparum*^48^ (Extended Data Fig. 5g, Supplementary Data 16). Finally, we performed GSEA separately at each timepoint to map time-specific stability changes (Fig. 5g, Supplementary Data 17). As expected, 20S proteasome, IF3 and IF4F maintained significant host-dependent stability differences throughout the entire IDC (Fig. 5h & i). Additionally, the ribosome was also stabilized in hRBCs at 23 hpi and remained stabilized until reinvasion (Fig. 5j), and mitochondrial respiratory complexes were also stabilized from 11hpi onwards hRBCs mainly (Fig. 5k).

## Discussion

### IDC in P. knowlesi, vivax and falciparum

In this study, we introduced a high-resolution transcriptional IDC time course of *P. knowlesi* cultured in human RBCs. Comparing this transcriptome to *P. vivax* IDC time course, we found only small differences between the two organisms, with significant gene expression peak differences of no more than 4 hours. This results diverges from a lower-resolution hRBC-cultured *P. knowlesi* IDC transcriptome recently published my Meulenaere et al^54^ who found that 25% of orthologous genes displayed expression peaks at different timepoints. We only identified 243 genes in our transcriptome did display larger expression shifts (>8 hours), these genes were not consistent in function. Importantly, we used GSEA to identify systematic shifts in functionally related groups of genes, whereas Meulenaere et al^54^ performed gene enrichment analysis (GEA), results of which can be influenced by a small number of genes without a systematic shift^55^. Notably, the difference in IDC rate between *P. knowlesi* and *P. vivax* was constant across the cycle. This is not the case for P*. falciparum*: the first 28 hours of *P. falciparum* development correspond to only 11 hours of *P. knowlesi* development, whereas the last 20 hours of both species progress at the same developmental rate. This marked difference in IDC pacing suggests that the trophozoite and schizont stages constitute the rate-limiting step of the IDC, likely owing to time-demanding processes including hemoglobin digestion, replication and preparation for invasion.

### *P. knowlesi* parasite face increased oxidative stress in human RBCs

Multi-omic profiling of *P. knowlesi* grown in two different host cells revealed major differences in enzymes involved in three closely related metabolic pathways. First, the GSS was expressed prematurely in hRBCs both on RNA and protein levels and displayed higher stability in ring stages when grown in hRBCs. Additionally, GR was more stable in hRBCs than in mRBCs throughout the entire IDC. Second, GS was stabilized and GDH was destabilized in hRBCs, indicating an increased metabolic flux from glutamate to glutamine rather than to 2-oxoglutarate and citric cycle. Third, the expression of PPP enzymes that produce NADPH was also shifted to earlier stages in hRBCs compared to mRBCs. These data show that in hRBCs, *P. knowlesi* (1) increases glutathione production and recycling, (2) switches metabolic flux towards the production of glutamine from glutamate, which is a precursor for glutathione, and (3) upregulates production of NADPH, which is used by GR for reduction of oxidised glutathione. This indicates an increased demand for glutathione and glutamine production in hRBCs.

In *P. falciparum*, inhibition of GS decreases asparagine levels^56^, consistent with the co-regulation of GS and AS observed here. Although culture media are supplemented with physiological concentrations of glutamine, inhibition of GS still impairs growth in *P. falciparum*, and knock-sideways of GS causes a 60% reduction in viability that cannot be rescued by increasing glutamine in the media^56^. The parasite’s remaining sources of glutamine are therefore de novo synthesis and hemoglobin digestion. We propose that the increased glutamine synthesis observed in hRBCs is driven by the most significant compositional difference between human and macaque hemoglobin: macaque hemoglobin α+β contains 7 glutamine residues, compared to only 4 in human hemoglobin, the largest single difference in amino acid composition between the two orthologs.

Glutathione is a key reducing agent that eliminates reactive oxygen species (ROS) in the highly oxidative environment of the RBC^57^. The early requirement for glutathione in hRBCs may likewise originate from differences in hemoglobin sequence: subtle sequence variation could render digestion less efficient, resulting in impaired hemozoin formation and increased ROS generation. Indeed, several proteases involved in hemoglobin digestion were more stable in hRBCs and mRBCs. In addition, basal glutathione concentration is significantly lower in hRBCs than in mRBCs^58^. It is therefore likely that the parasite draws on host-cell glutathione, which is depleted much earlier in hRBCs than in mRBCs, necessitating earlier engagement of parasite-driven synthesis. Moreover, *P. falciparum* has been shown to export glutathione into the host cell^59^. The increased demand for glutathione can therefore arise already in ring stages, before the initiation of hemoglobin digestion.

The premature expression of PPP proteins is likely linked to both glutamine and glutathione deficiency. PPP primarily supplies the NADPH required for glutathione recycling by GR, yet the metabolic pivot toward glutamine production simultaneously reduces flux through GDH, which is an alternative source of NADPH. This compounding demand for NADPH likely drives the observed upregulation of PPP. Notably, Plasmodium parasites lack transaldolase, which would otherwise metabolise erythrose-5-phosphate or sedoheptulose-7-phosphate into more common sugar phosphates^49^. Given the early expression of PPP proteins and lack of a complete shikimate pathway in *Plasmodium*^60^, we predict accumulation of erythrose-5-phosphate in ring-stage parasites in hRBCs, which would need to be metabolised later in the IDC.

Of note, the competing demand for glutamate to support both increased glutamine and glutathione production creates a metabolic bottleneck. Rewiring of these interconnected pathways, accompanied by changes in gene and protein expression timings of their component enzymes, likely reflects a key metabolic constraint the parasite must overcome to adapt to the host-cell environment.

### Dysregulation of protein homeostasis

Our data indicate that protein homeostasis in hRBC-resident *P. knowlesi* is disrupted at every step. First, protein expression of IF3 subunits, involved in translation initiation, was upregulated while their stability was decreased in parasite cultured in hRBCs compared to mRBCs. Second, the expression of the ribosomal proteins, especially those constituting the 40S subunit, increased at the RNA level but declined at the protein level in the late parasite stages cultured in hRBCs. Moreover, 18S rRNA levels dropped during the first half of the IDC in hRBCs. Third, proteins involved in protein folding, including the T-complex, showed increased but delayed expression in hRBC compared to mRBC-grown parasites. Finally, the 20S proteasome and HUB1 was both strongly stabilised and expressed early in hRBC-cultured parasites. In yeast, HUB1 regulates splicing under oxidative stress and plays a role in unfolded protein response^61^. We speculate that these changes are closely tied to the concurrent changes in glutathione and glutamine metabolism. As discussed above, oxidative stress is likely elevated as a consequence of low glutathione and glutamine levels in hRBCs. Although parasites eventually overcome this deficiency through increased glutathione production, our data suggest that GS is already saturated by 10 hpi, while protein levels do not effectively increase until 18 hpi. Meanwhile, *P. knowlesi*, with its compressed ring stage, begins digesting hemozoin as early as 8 hpi, imposing additional oxidative stress during early life stages. Increased ROS have been shown to affect ribosomes, including rRNA expression: superoxide treatment causes accumulation of 18S and 28S rRNA precursors^62^, ROS damage rRNA in *E. coli*^63^ and oxidise r-protein residues (reviewed in ^64^). Cysteine-containing r-proteins are particularly sensitive to mitochondria-derived ROS and have been proposed as key translational regulators under oxidative stress^65^. We have also previously found the *P. falciparum* ribosome to be indirectly linked to oxidative stress: artemisinin treatment, which induces ROS formation, increases the ribosome stability^46^. Moreover, glutamine deficiency may impose further stress on protein translation. Glutamine is a precursor for pyrimidine synthesis, an essential pathway in Plasmodium whose products are ultimately incorporated into mRNA^66^. While we observed increased metabolic flux toward glutamine and asparagine, the corresponding synthetases were neither upregulated nor temporally shifted, suggesting that the parasites cannot fully compensate for the glutamine deficit. Moreover, inhibition of GS in *P. falciparum* has been shown to induce phosphorylation of IF2α, which is linked to translation inhibition^56^. In human and mouse cells, charging of glutamine-tRNA is specifically sensitive to amino acid starvation, causing translational stalling^67^.

In summary, low glutathione levels in hRBCs, together with the reduced proportion of glutamine residues in human hemoglobin, induce oxidative stress that impacts translation, ribosome levels, 18S rRNA levels and protein homeostasis more broadly. With a delay, *P. knowlesi* compensates for these changes through increased ribosomal gene expression and metabolic rewiring. These adaptations represent potential barriers that *P. knowlesi* needs to overcome prior to production of gametocytes and successful human-to-human transmission.

## Supplementary figures

**Extended Data Figure 1.**
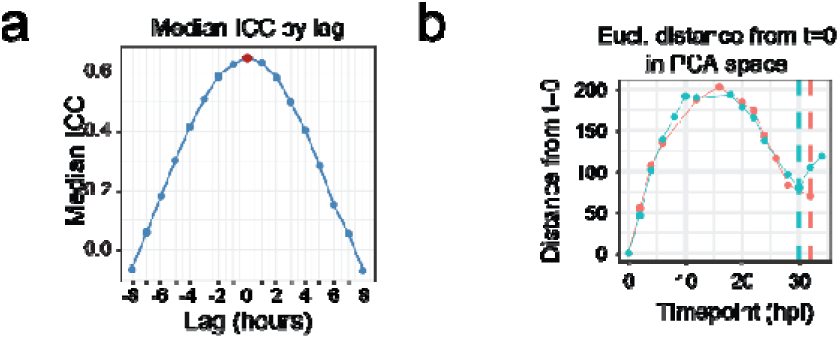
**a** Dependency of the ICC correlation between two replicates on potential lag shows that the two replicates are synchronous within 2-hour window. **b** Euclidean distances between timepoint 0 and other timepoints in the PCA space estimate the IDC length to 31 hours.

**Extended Data Figure 2.**
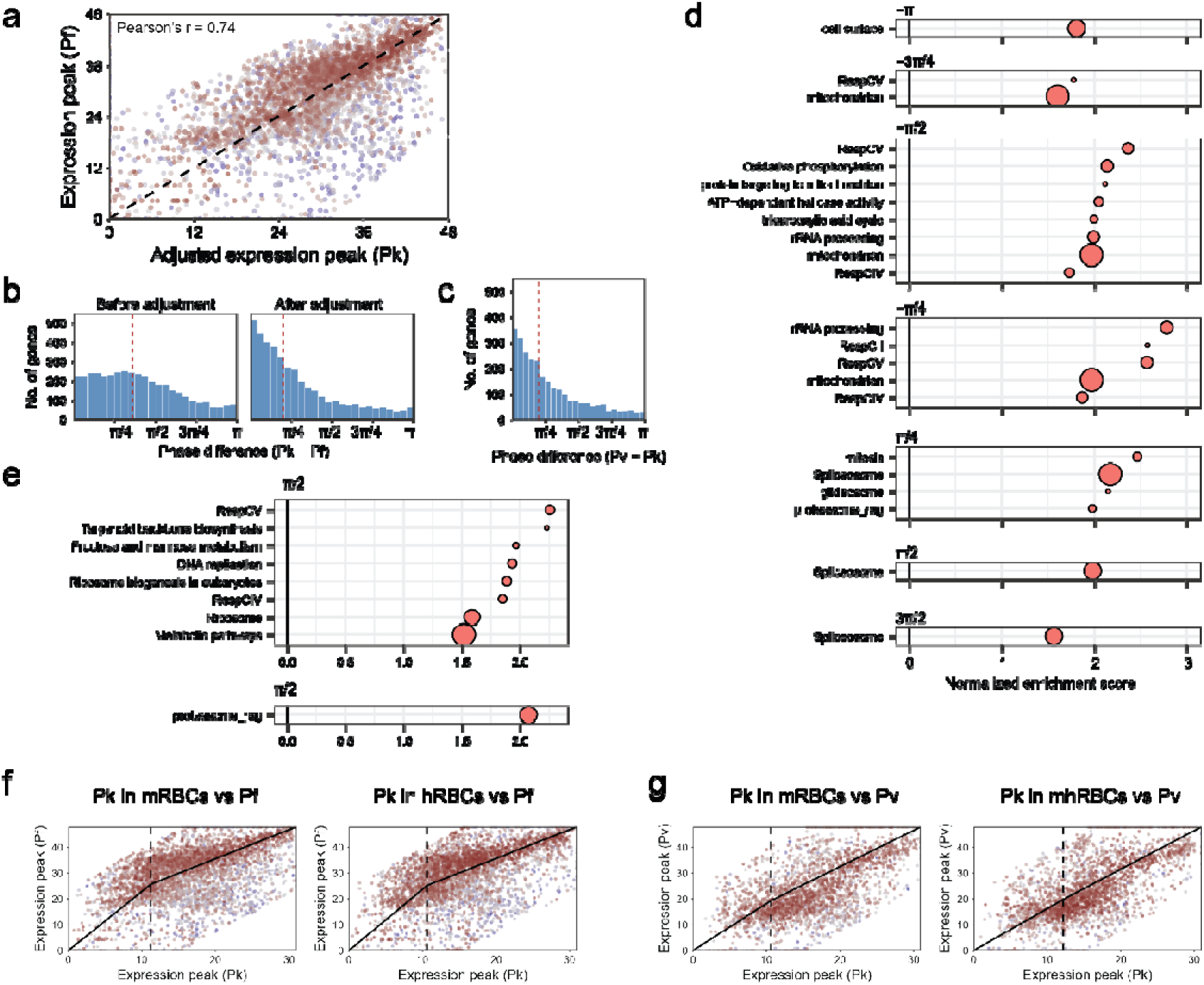
**a** Gene expression peaks show a linear correlation between *P. knowlesi* and *P. falciparum* after the adjustment of *P. falciparum* expression by two-segment function. **b** Absolute phase differences between genes of *P. knowlesi* and *P. falciparum* before and after adjustment. **c** Phase differences between genes of *P. knowlesi* and *P. vivax*. **d** One-sided GSEA across different gene expression time shifts between *P. knowlesi* and *P. falciparum*. **e** One-sided GSEA across different gene expression time shifts between *P. knowlesi* and *P. vivax*. **f-g** Correlation between expression peak times of *P. knowlesi* grown in macaque or human RBCs and *P. falciparum* (**f**) or *P. vivax* (**g**).

**Extended Data Figure 3.**
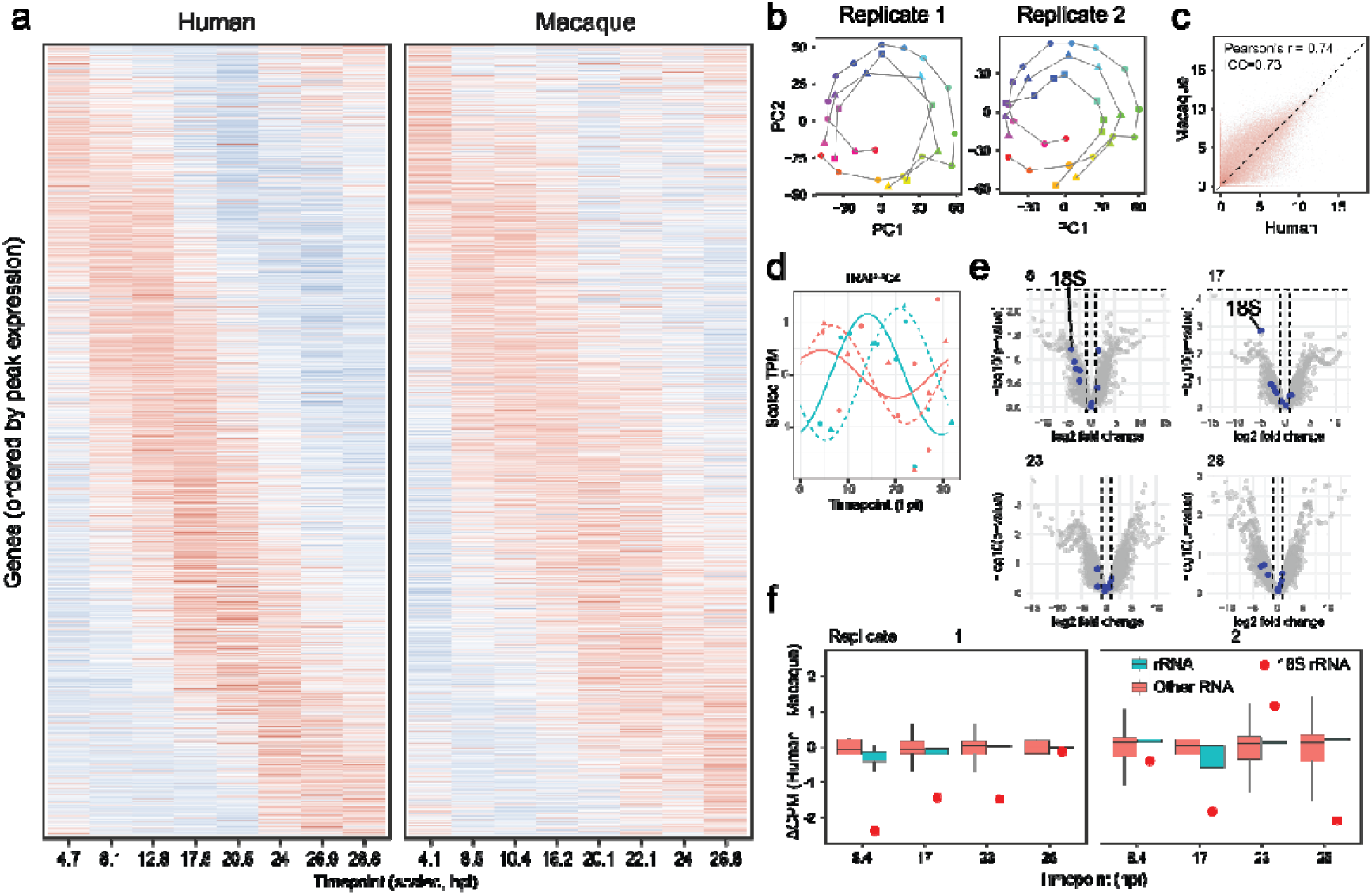
**a** Heatmap of scaled gene expression TPM values of *P. knowlesi* cultured in human or macaque blood, replicate 2. **b** PCA plots with human (square points) and macaque triangle points) overlayed on top of the consensus IDC (round points). **c** Correlation between log2 TPM values of *P. knowlesi* genes grown in human and macaque RBCs. **d** Expression profile of TRAPP complex subunit 4 in hRBCs and mRBCs. **e** Differential expression analysis at 4 timepoints. rRNAs are blue. **f** Difference in CPM (counts per million) values between hRBCs and mRBCs, comparing the distribution of rRNA and other RNA. 18S rRNA is highlighted by the red point.

**Extended Data Figure 4.**
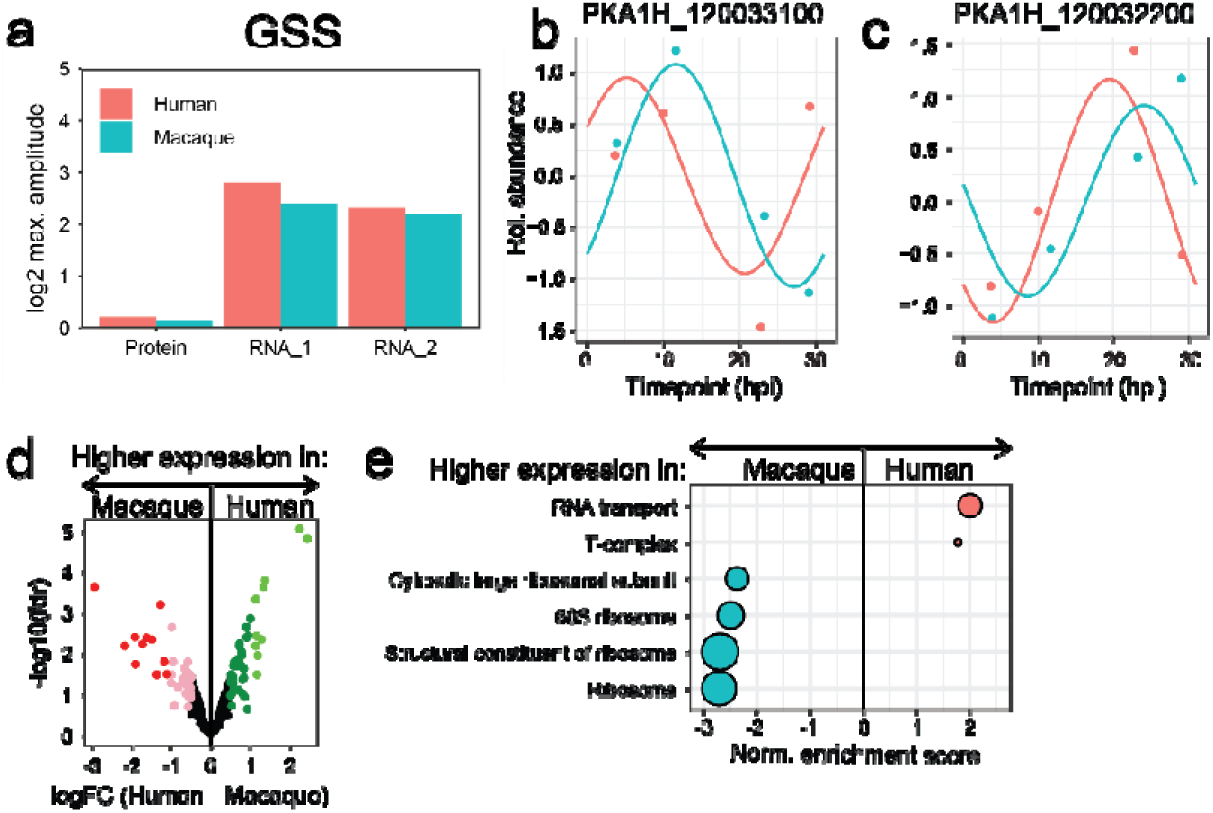
**a** Protein and RNA expression amplitudes of glutathione synthetase (GSS). **b-c** Gene expression profiles of glucose-6-phosphate dehydrogenase-6-phosphogluconolactonase (**b**) and 6-phosphogluconate dehydrogenase (**c**). **d** Differential expression analysis of proteins across the entire IDC. **e** Terms enriched by GSEA based on logFC values from the differential expression analysis.

**Extended Data Figure 5.**
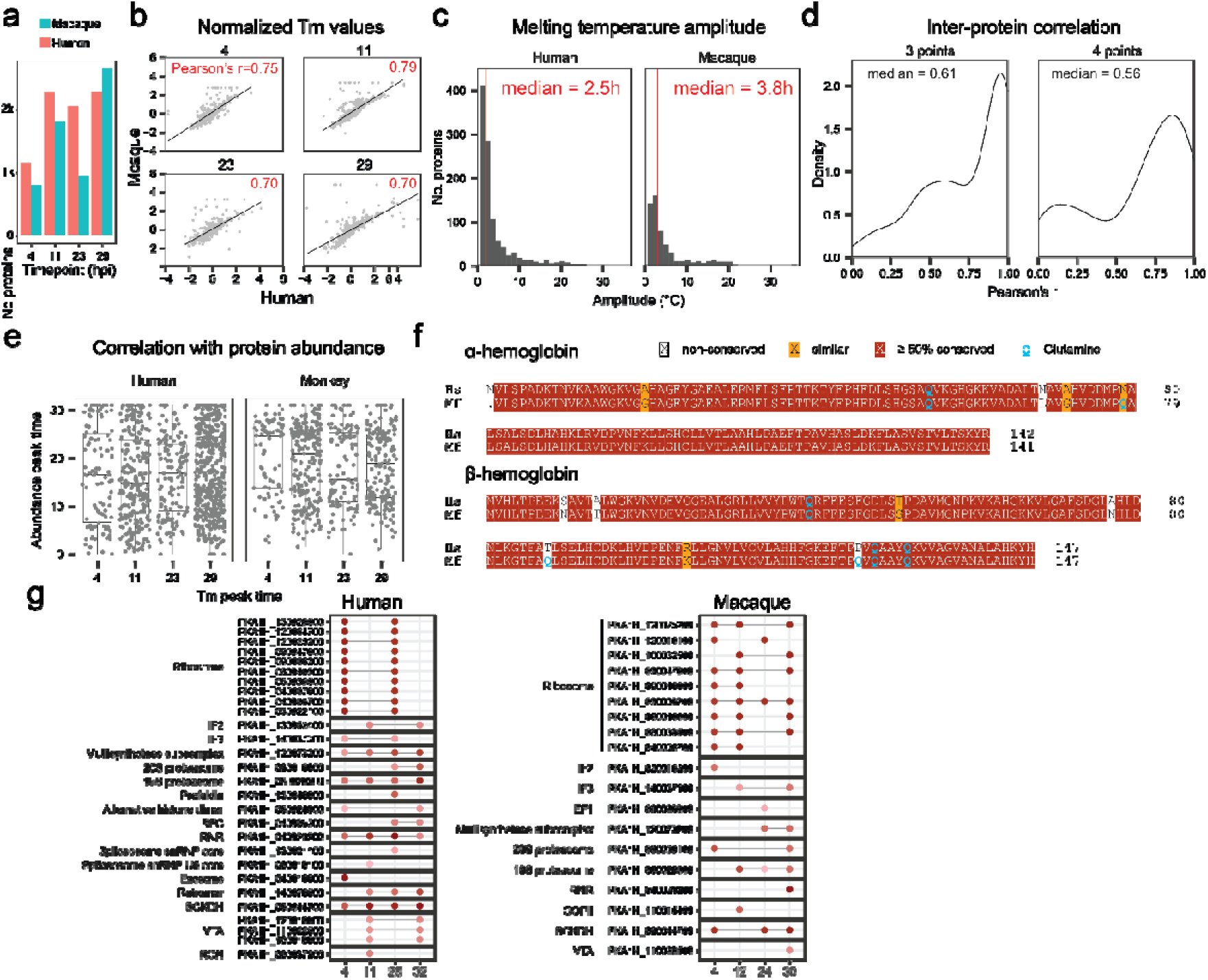
**a** Number of *P. knowlesi* proteins detected at different timepoints in macaque and human RBCs. **b** Correlation of melting temperatures (Tm) in hRBCs and mRBCs at different timepoints. **c** Amplitudes of melting temperatures of *P. knowlesi* cultured in human or monkey RBCs. **d** Correlation of stability profiles between pairs of proteins in hRBCs and mRBCs. **e** Distribution of abundance peak times at different stability peak times suggest that there is no correlation between the two. **f** Sequence alignment between human and macaque hemoglobins shows enrichment of glutamine residues in macaque hemoglobin. **g** Prediction of moonlighting proteins in human and macaque-cultured *P. knowlesi*.

## Resource availability

### Lead contact

Further information and requests for resources and reagents should be directed to and will be fulfilled by the lead contact, Zbynek Bozdech.

## Material availability

This study did not generate new unique reagents.

## Data and code availability

The sequencing data are deposited in the GEO repository under the accession number GSE348929. The mass spectrometry data are deposited in the PRIDE repository under the accession number PXD084821. The code used for the analysis of the data and the source data is available on Zenodo.

## Declaration of generative AI and AI-assisted technologies in the manuscript preparation process

During the preparation of this work, the authors used Claude Code (*Anthropic*) in order to streamline code writing for data analysis and Claude AI to streamline the Method section and organize the supplementary data. The authors reviewed and edited the code as necessary and take full responsibility for the content of the published article.

## Acknowledgements

We thank Rob Moon for providing us with *P. knowlesi* A1H1 strain. This work was funded by the Singapore Ministry of Education, grant # MOE2019-T3-1-007 and MOE-T2EP30223-0020, awarded to Zbynek Bozdech.

## Author contributions

Conceptualization: SP, NA, MK, ZB; Methodology: SP, NA, XG, AW, MK, PRR, ZB; Formal analysis: SP, KR, MK; Investigation: SP, NA, XG, AW, KR, ST, HD, KDG; Resources: PRR, ZB; Data Curation: SP, KR, HD, MK; Writing – Original Draft: SP; Writing – Review & Editing: SP, MK, PRR, ZB; Visualization: SP; Supervision: PRP, ZB; Funding acquisition: PRR, ZB.

## Methods

### Experimental Model and Study Participant Details

#### Parasite strain

*Plasmodium* knowlesi strain A1H1 was obtained from Robert Moon (London School of Hygiene & Tropical Medicine).

#### Animal subjects

Animal work was evaluated and approved by the National University of Singapore and SingHealth’s Institutional Animal Care and Use Committees (IACUC), protocol 2020/SHS/1609. Animals were housed following the Guide for the Care and Use of Laboratory Animals and the Association for the Assessment and Accreditation of Laboratory Animal Care (AAALAC) Standards. Long-tail macaque (*M. fascicularis*) blood was collected by venous puncture into EDTA-coated tubes (Vacutainer, BD). It was filtered using non-woven filters (Antoshin) to remove white blood cells and washed thrice with incomplete RPMI 1640 medium (iRPMI, Gibco) and stored at 4°C .

#### Human subjects

For both routine parasite maintenance and experiments, human blood (Duffy-antigen positive) was obtained from consenting volunteers recruited following the guidelines of the Nanyang Technological University’s Institutional Review Board (protocol IRB-2021-1015) under Singapore’s Human Biomedical Research Act 2015 rev 2020. Samples were anonymized on extraction, centrifuged to remove plasma and buffy coat, washed twice with iRPMI and stored in iRPMI at 4°C for up to two weeks.

## Method Details

### Parasite culture

*Plasmodium knowlesi* (strain A1H1) was cultured using standard Pk medium, composed of RPMI 1640 with 25 mM HEPES (Sigma) supplemented with Albumax I (Gibco) (5 g/L), hypoxanthine (Sigma) (0.3 mM), dextrose (Sigma) (4 g/L), L-glutamine (Gibco) (40 mM), gentamicin (Gibco) (10 mg/L) and heat-inactivated horse serum (Gibco) (10% v/v). Cultures were kept static in P75 vented flasks (Corning) at 2% haematocrit in a gassed incubator (Esco) at 37°C with 5% CO_2_, 3% O2, and 92% N_2_. Culture medium was changed daily. Parasitaemia and parasite morphology of the cultures were assessed by optical microscopy observation of Giemsa-stained (Sigma) blood smears. Fresh packed RBC (either macaque or human, depending on the culture) were added as needed to adjust the parasitaemia to between 1-5% for routine culture, or up to 15% before synchronization to start a lifecycle experiment. For parasite synchronisation, gradient centrifugation was used on cultures at 50% haematocrit on a layer of Nycodenz (Histodenz, Sigma), to enrich for schizonts to >85%.

### Transcriptomics sample collection

After synchronization, the parasites grown in human or macaque RBCs were recovered for one IDC and after 90% of schizonts invaded, parasitaemia was adjusted to 1% and the culture was sampled every two or six hours. At sample collection, the RBCs were lysed in 8x vol of TriZol and kept at -80°C until RNA extraction. Total RNA was extracted using Quick-RNA Miniprep columns (Zymo Research) following manufacturer’s instructions. Libraries were prepared using Zymo-Seq RiboFree® Total RNA Library Kit (Zymo Research), with 100 µg RNA, 4 h incubation for rRNA-depletion and 14 amplification cycles. Molarity of libraries was determined using qPCR; libraries were pooled equimolarly and sequenced on Novaseq S4 instrument (Illumina) with pair-end 150 bp long reads protocol, generating on average 3-4 Gb of data per sample (NovogeneAIT Genomics Singapore Pte Ltd).

### RNA-seq read processing and transcriptome quantification

Sequencing reads were quality- and adapter-trimmed with Trim Galore (Cutadapt-based) in two passes: the first pass removed library-specific dual-index adapters (Q30, minimum retained length 75 bp), the second removed residual generic Illumina adapter sequence (minimum retained length 35 bp). High-quality reads were aligned to the P. knowlesi A1H1 reference genome using HISAT2 (--dta, --min-intronlen 20, --max-intronlen 3000, --rna-strandness RF), and alignments with more than three reported positions (NH tag) were discarded before sorting to BAM with samtools. For each sample, filtered alignments were assembled in strand-specific mode with StringTie; per-sample assemblies were merged into one consensus transcript set, and each sample was then re-quantified against this merged annotation using StringTie’s reference-guided, estimate-only mode (-e). Raw per-transcript read counts were extracted from the StringTie coverage output with the prepDE.py3 companion script.

Before normalization, the count matrix was filtered using custom script in R. rRNA transcripts and SICAvar (type I/II) and KIR variant-antigen transcripts were removed by annotation; samples with fewer than 0.8 million reads remaining after this gene-level filtering and samples with low within-timepoint correlation that also had low neighbouring-timepoint correlation (r<0.7) were excluded. TPM and FPKM values were then calculated from the filtered counts by length-normalization. For temporal IDC profiling, TPM values were log2(x+1)-transformed and then z-scaled per gene within each replicate. The length of the IDC was estimated from the scaled-data PCA: for each replicate, the Euclidean distance in the space of the first 10 principal components between every timepoint and the t=0 sample was calculated, and the timepoint (beyond a 10 h minimum offset) with the smallest distance from t=0 was taken as the completion of one cycle. The estimate was averaged across replicates. Agreement between the two replicates was assessed in three ways: (i) the intraclass correlation coefficient (ICC, two-way, agreement, single measures) between the scaled expression matrices; (ii) a lag-scan, in which ICC was recomputed after shifting replicate 2 by −8 to +8 h in 1 h steps (modulo the estimated IDC length) to confirm the replicates were not offset in time; and (iii) the Pearson correlation between log2 TPM values of the two replicates, and its dependence on transcript abundance. For each gene and replicate, the scaled expression time course was fitted with a cosine function *y* = 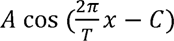 by non-linear least squares (Levenberg–Marquardt algorithm, R package *minpack.lm*^68^), with period T fixed at the estimated IDC length; the fitted phase C was taken as the peak expression time and the fraction of variance explained (R²) as a goodness-of-fit measure. Agreement between replicates’ phases was quantified as the circular phase difference and by the circular correlation coefficient. The consensus transcriptome across the two replicates was built gene-by-gene: where a gene’s phase estimate was available in only one replicate, that replicate’s values were used; where both were available and their phase difference was < π/4 (good agreement), the mean TPM across replicates was used; otherwise, the replicate with the higher R² was used. The consensus time course was then refitted to obtain final consensus phases and R² values, which define gene order and peak-expression timing in all subsequent analyses. Timepoints from subsequent host-comparison experiments were re-mapped onto the consensus IDC timescale based on combined scaled-data-based PCA, on which smoothing spline was applied to consensus trajectory in the first 5 principal components and each human- or macaque-host sample’s timepoint was re-assigned to the value on that spline trajectory closest to its own PCA coordinates. For each gene, the circular phase difference (human − macaque) was averaged (circular mean) across the two replicates after restricting to fits with R² ≥ 0.5; a gene was called host-differential if the phase difference exceeded π/4 in both replicates with the same sign.

### Comparison of IDC pacing with *P. falciparum* and *P. vivax*

Published *P. falciparum* IDC transcriptome data^37^ and *P. vivax ex vivo* transcriptome data (three strains; the smru1 series was used^40^) were each log2-transformed, gene-wise z-scaled and fitted as described above, using each species’ own reference cycle length (48 h). Orthologous genes were mapped between species using ortholog table available from malaria cell atlas^69^. Circular correlation between orthologs’ peak phases (restricted to genes fitted with R² > 0.5 in both species) was used as an overall measure of temporal concordance. To characterize how the pace of IDC progression differs between species, peak-expression scatter plots (ortholog phase in *P. knowlesi* vs the comparator species) were fitted with a two-segment, piecewise-linear model: a breakpoint and a shared join value were optimized by grid search to maximize the orthogonal-distance R² (weighted by the mean R² of each ortholog pair) summed across the two segments. As an independent check, the circular phase difference between species (comparator minus *P. knowlesi*, expressed in hours on the comparator’s timescale) was modelled as a function of the *P. knowlesi* phase using a saturating exponential fitted by non-linear least squares.

### Differential gene expression analysis

Raw gene counts (samples were matched to four common timepoints of approximately 8, 17, 23 and 28 hpi, ±2 h tolerance) were compared between hosts using edgeR: for each timepoint, a DGEList was built from the raw counts, library sizes were TMM-normalized, gene-wise dispersions were estimated with a robust empirical Bayes shrinkage towards a common trend from a design blocking on the 2 h/6 h experiment as a paired replicate factor and host effects were tested by a generalized linear model quasi-likelihood F-test on the host coefficient. Differential logFC values from each timepoint were tested by GSEA and ribosomal (40S/60S) and respiratory-complex (III/IV/V) subunit logFCs were compared to all other genes by Wilcoxon rank-sum test at each timepoint (Benjamini-Hochberg adjusted p-values).

### Proteomics sample preparation, measurement and data processing

Parasites cultured in human or macaque RBCs were synchronized as described above and cultured for one IDC to reach 30% parasitaemia. The trophozoite and schizont stage parasites were additionally enriched by magnetic sorting and the intact cell meltcurves were generated across ten temperature points as previously published^48^. The cells were mechanically lysed, solubilized in 1% NP-40 and the soluble fractions were recovered. The peptides were generated using the standard processing protocol^70^ and 10 sample from each meltcurve were TMT-labelled, desalted and fractionated as previously described^48^ for stability proteomics measurements. For quantitative proteomics, the cultures were prepared in the same way, and the samples were collected without thermal denaturation. Mass spectrometry measurements were conducted on Dionex 3000 UHPLC system coupled with a Q Exactive HF mass spectrometer (Thermo Fisher Scientific). Peptides were separated on 50cmx75μm analytical column (EASY-Spray™, Thermo Scientific) over a 70-minute gradient (1-55 min: 2-25%, 55-57 min: 25-50%, 57-58 min: 50-85%, 58-63 min: 85%, 63-70 min: 2%). Mass spectrometric data were acquired using data-dependent acquisition (DDA) with full scan MS spectra (350-1550 m/z, resolution: 70,000, AGC target: 3e6) and Top12 MS2 spectra (resolution: 35,000, AGC target: 1e5, 1e5 isolation window at 1.2 m/z). Raw data were analysed using Proteome Discoverer 2.1. The raw spectra were searched using Sequest HT against P. falciparum 3D7 protein sequences translated from the Sanger Institute Coding Sequence database (plasmoDB release 52) and the Uniprot human protein database (accessed in April 2021). The search parameters used were as follows: MS precursor mass tolerance of 25 ppm, fragment mass tolerance 0.05 Da, maximum 3 mixed cleavages, static modifications: carbamidomethyl, TMT10plex; dynamic modifications: oxidation (M), deamidated (NQ), acetyl (protein N terminus). Forward/decoy searches were used to estimate FDR. Peptides and peptide spectrum matches with high and medium FDR levels (up to 5%) were accepted.

### Differential protein expression analysis

Protein abundances were variance-stabilised and normalised across samples using the vsn package . For differential expression analysis, abundances were pooled across timepoints within a host and compared between hosts using a limma-based moderated linear model (design matrix on host, with timepoint as a blocking factor), with hits called at FDR < 0.05 and |log2FC| ≥ 1 (candidates at FDR < 0.25, |log2FC| ≥ 0.5).

### Thermal proteome profiling analysis

Meltcurve data were scaled, normalized and fitted with sigmoidal models using *MAPX* R package^48^ to calculate melting temperatures for each protein (Tm). Tm values were robustly z-scaled (median/MAD) within each host/timepoint, then again across timepoints within each host. Normalised Ti was correlated between hosts (Pearson, overall and within timepoint-matched subsets), and each protein’s stability-profile amplitude (peak-to-trough) was compared between hosts. For each protein and timepoint, ΔTm was defined as the normalized Tm difference (human − macaque). The differential stability analysis was performed using limma-based moderated linear model^72^ as described above (hits at FDR < 0.05, |log2FC| ≥ 1; candidates at FDR < 0.2, |log2FC| ≥ 0.5). ΔTm time course of individual proteins was fitted with the cosine function to visualize the timing of the largest host-dependent stability shifts. Selected protein complexes were each compared to all other proteins by Wilcoxon rank-sum test at each timepoint (Benjamini-Hochberg-adjusted p-values).

### Transcriptome vs proteome comparisons

The protein abundance profiles were z-scaled and cosine-fitted as described above. Phases extracted from the cosine fits were subtracted (protein phase – RNA phase) to calculate the translational lag. The distribution of these lags were summarized per host by circular kernel density estimation and described by circular median. The amplitudes were calculated as the difference between maximum and minimum log-scaled gene expression or protein abundance values.

### Gene set enrichment analyses

Gene sets were drawn from the Kyoto Encyclopedia of Genes and Genomes (KEGG), gene ontologies (GO) and in-house-curated protein complexes^48^ and the enrichments were calculated using R package *fgsea* based on logFC values calculated as described above or ΔTm values for timepoint-by-timepoint GSEA for stability proteomics. For peak expression time-based GSEA, a series of reference timepoints spanning the IDC in 2h (consensus transcriptome) or 1/8 of the IDC (human vs macaque comparisons) steps, each gene’s cosine-fitted phase was converted into a peak-proximity score (maximal at the reference timepoint, decaying with circular distance from it, folded at half the period), and this score was used to rank genes for GSEA. Highly overlapping enriched gene sets were always collapsed.

### Moonlighting profiling

Meltcurve-derived features were reduced by PCA analysis using *MAPX* R-package^48^ and for each of complex from the curated list of protein complexes, subunits were partitioned into two clusters by k-means (k = 2, 100 iterations/starts) in the PC space (weighted by the variance explained by each PC). A subunit was called a candidate “moonlighter” when it consistently significantly separated from the bulk of its complex across timepoints (present in ≥ 2 timepoints), identifying subunits whose thermal behavior differs from the rest of their complex.

### Hemoglobin amino-acid composition analysis

Human and Macaca fascicularis hemoglobin alpha- and beta-chain sequences were retrieved from UniProt (accessions P69905, P68871, P21767 and P68223) via the UniProt REST API and parsed with Biostrings. Amino acid composition was counted per chain, summed across the alpha and beta chains to give a total hemoglobin amino acid composition per species, and expressed as a human-to-macaque count ratio per residue to identify amino acids over- or under-represented in the human protein. The alpha- and beta-chain sequences were separately aligned between species to compare residue-level conservation, using the Bioconductor msa package, with alignments rendered via TeXshade.

